# The Relationship Between Soldier Height, Weight and Army Combat Fitness Scores

**DOI:** 10.1101/2021.12.22.473875

**Authors:** Ryan Palmer, Mark DeBeliso

## Abstract

The United States Army recently implemented the Army Combat Fitness Test (ACFT) which was designed to more accurately measure functional-combat fitness constructs. The ACFT replaced the former Army Physical Fitness Test (APFT). The three advent APFT consisted of: two-minute push-ups (PU), two-minute sit-ups (SU), and a timed two-mile run (RUN). The ACFT consists of six events; 3 Rep Max Deadlift (MDL), Standing Power Throw (SPT), Hand Release Push-up (HRP), Sprint-Drag-Carry (SDC), Hanging Leg Tuck (LTK), and a timed two-mile run (2MR). This study investigated the relationship between Soldier height (1.79±0.07 m) and weight (body mass 86.8±14.2 kg, BMI 27.1±3.9) on ACFT scores (442.3±54.4) of 655 male U.S. Army National Guard Soldiers in a Field Artillery Brigade. For the purpose of the investigation body mass index (BMI) was calculated as the metric representing the Soldier’s height and weight. The mean and standard deviation (sd) were calculated for the ACFT event and total scores. Pearson correlation coefficients (PCCs or r) were calculated between BMI and ACFT event and total scores. Likewise, PCCs were calculated between the ACFT event and total scores. The ACFT mean±sd scores were as follows: MDL=92.2±31.8 (3 maximum repetitions), SPT=9.5±2.2 (meters), HRP=24.6±13.1 (repetitions), SDC=119.8±21.7 (seconds), LTK=6.2±5.4 (repetitions), 2MR=1095.0±233.7 (seconds), ACFT total score=442.3±54.4 (points). Significant positive correlations were found between the ACFT total score: MDL (r=0.70), SPT (r=0.50), HRP (r=0.74), and LTK (r=0.76) events. Conversely, significant negative correlations were identified between ACFT total score: SDC (r=−0.68) and 2MR (r=−0.53) events. Within the parameters of this study, Soldier BMI demonstrated “no to weak” association with individual ACFT event or ACFT total scores. Further, the range of PCCs between the ACFT event scores were “no to moderately high”. Military leaders may consider the results provided as combat and fitness tests continue to evolve.

## INTRODUCTION

From the earliest days at West Point U.S. Military Academy, Soldiers have been physically tested in numerous ways including; obstacle courses, ruck marches, horse rides, swims, rope and wall climbs, jumping a ditch, jumping over a horse, dashes of various distances, as well as other physically demanding activities (8,34). These physical assessments were constructed to measure a Soldier’s ability to perform soldiering skills required for combat, much like an athlete specifically trains for a specific sport. Unlike sports, the requirements to be successful in combat change with every theatre of conflict.

The Army Physical Fitness Test (APFT) was developed in 1980 and became the Army standard in 1982 (20). The APFT consisted of timed push-ups, sit-ups and 2-mile run which primarily measured muscular endurance (15,17) and could be performed almost anywhere without equipment. Recognizing that modern combat requires a much greater physical demand on Soldiers, the U.S. Army Training and Doctrine Command (TRADOC) has released a new physical training program designed to meet the needs of the modern Army to have strong, agile, fit Soldiers and at decreased risk of injury (15,22).

The Army Combat Fitness Test (ACFT) was developed to replace the 3-event APFT (14,37). The ACFT is a 6-event combat fitness test based on high intensity functional movement exercises while maintaining the endurance run. The ACFT events are graded as separate tasks scored individually where the sum of the six tasks determines a composite score for the test. A minimum standard of 60 points must be met in each event to constitute passing ACFT, regardless of total score. The events are in order: Three repetition max dead lift (MDL), standing power throw (STP), hand release push-ups (HRP), sprint-drag-carry shuttle (SDC), hanging leg tuck (LTK) and a two-mile run (2MR).

Modern day Soldiers are required to carry a minimum of a 35 lb. (15.87 kg) vest, 8 lb. (3.62 kg) rifle, 3 lb. (1.36 kg) helmet and up to an additional 100 lbs. (45.35kg) of gear. Ruck marching remains a staple of the Army and is often conducted at distances between 3 and 18 miles (7,39). In the present urban combat environment, Soldier movement patterns require high anaerobic demands of agility, acceleration, and moving to and from a prone, or kneeling, position while loaded with gear. These physical demands require Soldiers to not only be strong but also powerful, agile and have the stamina to endure (15). Several items were taken into consideration for the implementation the ACFT (37). Does it prepare Soldiers for movement patterns that they will perform in combat situations? What is the risk for injury with a new program? Can measures be put in place to reduce the risk of injury? Does it decrease body fat, preserving the image and associated health benefits from lower Body Mass Index (BMI)?

Aerobic exercise has been a gold standard for military physical fitness and maintaining healthy body weight (6). Soldiers formerly trained to pass the APFT with high volume endurance exercises; however, several authors have found that this type of training is not effective in preparing Service Members for combat (25,27,28). The modern ACFT and the associated TRADOC training program is structured using high-intensity functional training (HIFT) as it is believed to be more specific to combat performance and other military tasks (3,4,5,18,21,23,29,30,31,33,36,40,41). Studies show that HIFT does not negatively affect performance on the two-mile endurance run and some studies support that run performance is increased following HIFT training (21). High intensity functional training has also been shown to have a positive effect on BMI (3,24,30). Injury occurrence of the ACFT and TRADOC training program appears to be similar to the former APFT and military preparatory training (9,10,24).

Although the Army’s physical assessment has changed, the body composition standard has not, but rather become a metric of military physical fitness that is heavily debated among Soldiers. Currently, the Army’s height and weight requirement is designed to keep Soldier’s body weight low and composition lean. Research indicates that healthy individuals have a lower BMI which is traditionally determined using a height to waist circumference calculation (38).

The Army Body Composition Program is structured on a height and weight scale where any Soldier who exceeds the weight allowance for their height is required to ‘tape’. A circumferential measurement of the neck and waist (neck, waist and hips for female Soldiers) is used to calculate a Soldier’s percent body fat. The average of three neck measurements are subtracted from the average of three waist measurements and recorded as the circumference value. Using the Soldiers height and circumference value, the estimated percent body fat is determined from a table in Appendix B of Army Regulation 600-9 (12). For this study, body composition will be referenced as BMI and Army body composition as height/weight (HT/WT). It is acknowledged that both of the aforementioned references to body composition are estimates with known limitations.

Because Army body composition estimates percent body fat from neck to waist circumference, many argue that the system is inaccurate and favors fatter Soldiers over muscular Soldiers. A study by Babcock et al. (2) supports that the Army circumference scale is inaccurate when compared to skin-fold measurement. Another argument is; the physical demands of combat require larger framed Soldiers. Some research supports that Solders who have a larger waist (BMI>25) are more successful performing functional movement under load, especially load carriage (32). In support, Grier et al. (11) determined that physically active young men may be misclassified by BMI by not differentiating between lean and fat body mass. This may be important information for military Commanders to consider as part of their force structure.

### Problem Statement

Athletes are becoming, bigger, faster and stronger to stay competitive. Likewise, Soldiers are getting larger and stronger in order to meet the demands of their duty. A recent study by Keefer & DeBeliso (17) examined the relationship between the US Marine Physical Fitness Test scores and US Marine Combat Test scores yielding no-moderate correlations. The study did not however assess the relationship of body composition to physical assessments (17). Now that the Army has transitioned from the APFT (aerobic endurance) to the ACFT (HIFT and aerobic endurance), a few questions need to be asked. Does a relationship exist between BMI and ACFT performance? Specifically, what is the relationship between BMI and individual ACFT event scores? And, what is the relationship between BMI and the ACFT total score?

Due to the lack of published research and to the best of our knowledge, this is the first study to explore the relationship between Soldier HT/WT and ACFT performance.

## METHODS

### Participants

Data was collected from U.S. Army Soldiers in the 65^th^ Field Artillery Brigade, a combat military occupation specialty (MOS). The ACFT has tiered scoring where combat MOS’s are expected to perform the highest minimum standard. All participants were familiar with, and practiced on, the events prior to being tested. Scores are independent from gender and age. Data redacted of personal identifiable information (PII) was collected to conduct the study. Researchers were not present at the testing locations. Permission from the Institutional Review Board and the U.S. Army was obtained prior to obtaining and analyzing the data.

The participant sample examined in the current study was comprised of 655 male, Officer and enlisted United States Army National Guard Soldiers. The data examined was recorded by, the individual military unit administrator, and entered into the Digital Training Management System (DTMS). This is the Army’s official record for fitness testing and height and weight records. Data was collected during the period of between 06/30/2021 to 09/30/2021.

Permission to conduct this study of existing data was obtained from the Institutional Review Board at Southern Utah University and was approved as an exempt status (SUU IRB approval #*18-032021a*). Approval was also obtained from the Army Research Protection Administration Review Board on memorandum signed 07 June 2021 by the Research Ethics and Compliance Officer, in accordance with DoDI 3216.02. “This research was carried out fully in accordance to the ethical standards of the International Journal of Exercise Science” as described previously (26).

G*POWER 3.1.9.2 (Universitat Kiel, Germany) software was utilized to conduct a power analysis which indicated that 44 participants were required. The power analysis was based on the following assumptions: medium-high effect size of ES=0.40, statistical power 1-β=0.80 (two-tailed), and α=0.05. Sample size examined in the current study consisted of n=655 National Guard Soldiers.

### Protocol

Data for this study was obtained from the U.S. Army Digital Training Management System (DTMS). The ACFT data consisted of individual event scores: MDL weight, SPT distance, HRP repetitions, SDC time, LTK repetitions and 2MR time. Anthropometric data consists of height and weight for all Soldiers; as well as neck and waist measurements for Soldiers who exceeded the Army HT/WT standards. Sex was included in the data as Army body composition standards are different for male and female Soldiers. Because of the differing measurement standards, female data was removed from this study. All data was redacted of personal identifiable information prior to being sent to the researchers.

Testing was conducted by trained evaluators following current Army Regulation FM 7-22 and the ACFT Field Manual (13,14) (Figure 1). Soldiers had a total of 70 minutes to complete the ACFT. Each event was demonstrated, scored and recorded by a qualified evaluator on a Department of the Army (DA) Form 705 (13). Measurement units were: lbs. (MDL), meters (SPT), repetitions (HRP), time (SDC), repetitions (LTK) and time (2MR) respectively. For statistical reporting, measurements were converted to the metric equivalent. Raw scores were converted to points using the ACFT standards scale (14). Scores are not specific to gender or age. Each event was conducted and scored as follows:

**Figure 1.**
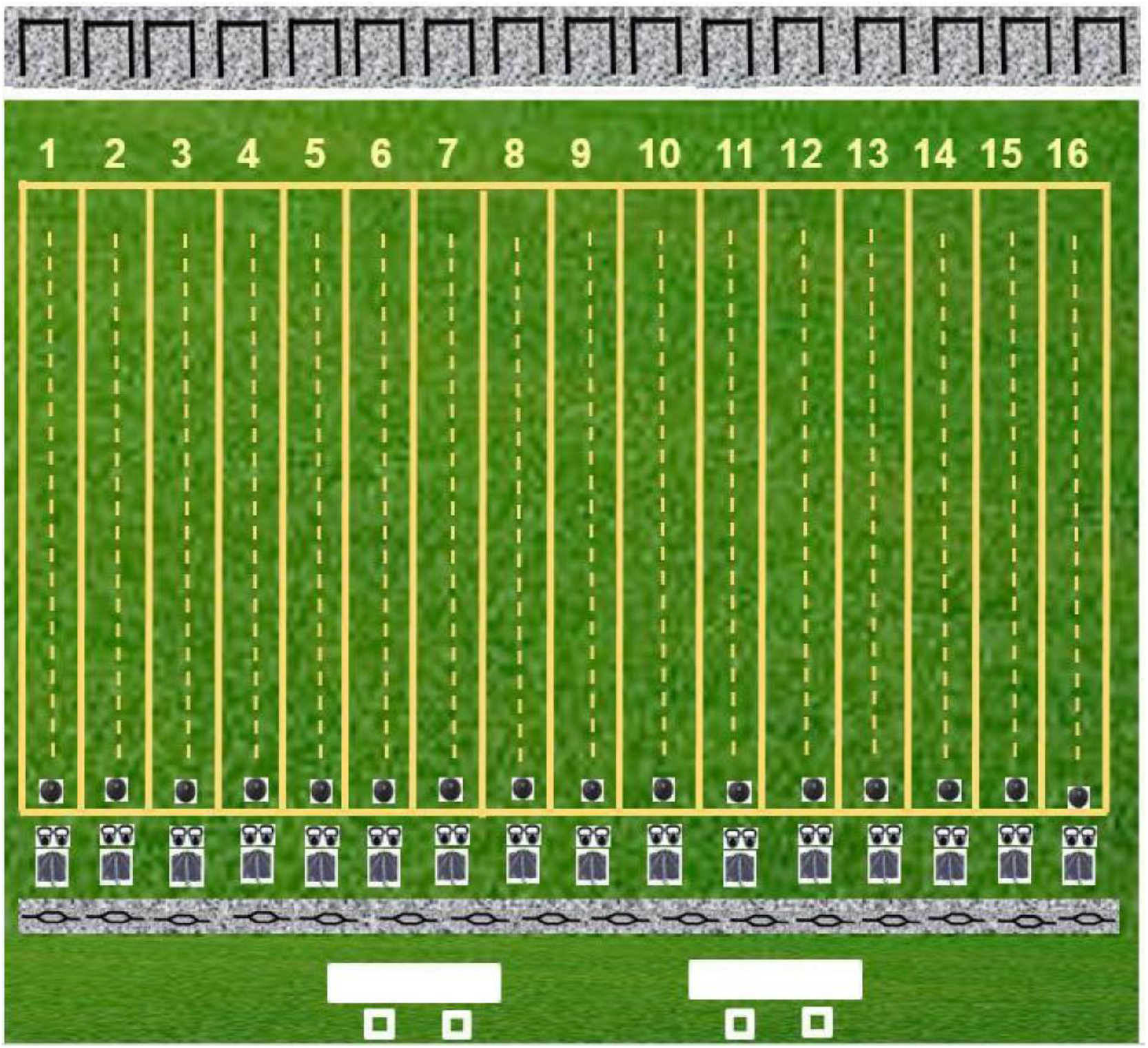
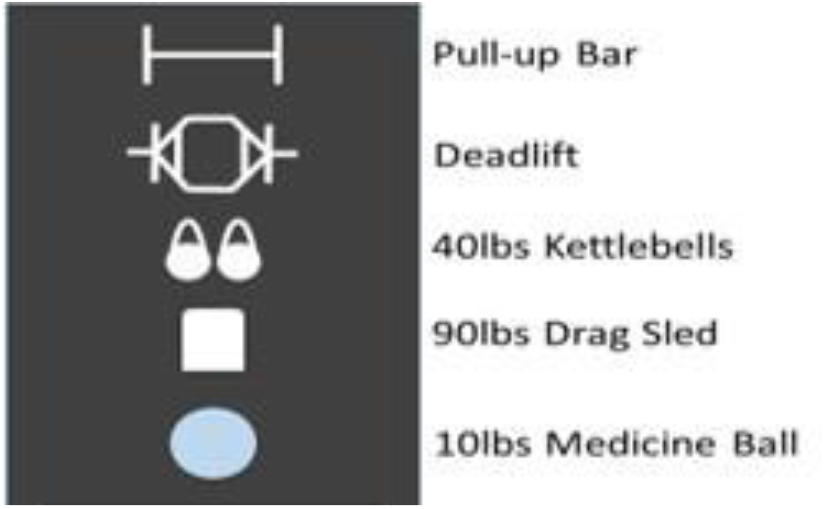
ACFT lane layout. Diagram not to scale. All six events occur within the same lane (25 m long by 3 meters wide). (Figure reprinted with permission of the U.S. Army). (14)

The three repetition max dead lift (MDL) is divided into three phases. First is the preparatory phase where the Soldier steps inside an Olympic hex bar, and takes a proper grip and stance. The second phase is the upward movement, or pull. During this stage, the Soldier’s hips were not permitted to raise above or before the shoulders. Their back was to remain straight through the pull. The Soldier slightly paused movement at the top of the pull before beginning the third phase. In the downward phase, the Soldier lowered the weight by flexing the hips and knees, but not the back. Weight was lowered until the plates touched the ground. No dropping or bouncing the weight was allowed. Soldiers performed three consecutive repetitions. The heaviest weight successfully lifted was recorded as a raw score. The event was stopped for safety if the hips moved above shoulders, there was excess rounding of the shoulders, genu varus with the pull or if the Soldier dropped the weight (13,14).

The standing power throw (SPT) is commonly known as a backward overhead medicine ball throw. The event began after a maximum two-minute rest from the previous event. Soldiers performed the throw using a rubberized 10 lb. (4.5 kg) medicine ball. The Soldier begins the throw by holding the ball in both hands while squatting with the ball between their legs in a loading phase. Next the Soldier extends their legs, hips, trunk, shoulders and arms, releasing the ball backwards overhead. Each Soldier performed one practice throw, followed by two throws for record. The thrower cannot step on or over the throwing line. The spot where the ball lands was marked by a grader and measured in meters. The throw achieving the greatest distance was recorded for points. The practice throw was not recorded. Safety precautions included: ensuring that the ball clean and dry and graders were in a safe location prior to throwing (13,14).

The hand release push-up (HRP) was started after a maximum three-minute rest from the previous event. Soldiers began in a prone position with their hands flat on the ground and index fingers aligned below the lateral edge of their shoulders. The Soldier’s chest, hips, thighs and toes were in contact with the ground. The first phase is the push. The Soldier raised their body as a unit, maintaining a straight line from shoulders to ankles, until elbows were locked at full extension. In the second phase, the Soldier lowered their body as a unit, returning to the prone position. Unlike a traditional hand release push-up, where hands are raised from the ground between each repetition; phase three of the HRP required Soldiers to fully extend their arms laterally from the chest to 90 degrees of shoulder abduction. Once fully extended Soldiers returned their arms to the starting position to begin the next repetition. Their hands were permitted to stay in contact with, or raise from the ground while extending the laterally. Only correctly performed repetitions were counted. The maximum number of correctly performed push-ups was recorded as a raw score. A Soldier could only rest in the up position. Failure to push up after five seconds on the ground or lifting any hand or foot off the ground, terminated the event (13,14).

The sprint-drag-carry (SDC) shuttle was started after a maximum three-minute rest from the previous event. Soldiers began the event from a prone position behind the starting line. After the signal to “Go”, the Soldier would jump to their feet and sprint 25 meters, where they had to touch the mid-line with the foot and one hand before sprinting back to the starting point. The second leg of the shuttle is a 90 lb. (40.8 kg) weighted sled drag. The sled was drug by back peddling until the entire sled crossed the line on either end of the 25-meter course. Jerking on the straps, potentially causing shoulder or back injury, was not allowed. The third leg is a lateral shuffle and the Soldier were to remain facing the same direction, down and back, as to lead with opposite legs. At the mid and end line, the Soldier touched the line with both foot and hand. The event was stopped for safety if the Soldier’s legs cross over one another. The fourth leg of the SDC is a weighted carry of 40 lbs. (18.1 kg) kettle bells, one in each hand. Soldiers only touched the mid and end line with the foot in this phase. For safety, Soldiers changed direction under control, and were not permitted to throw or drop kettle bells at the finish line. The final sprint was conducted as the first, but the Soldier started in a standing position, and ran through the end line. Time was recorded to the nearest whole second as a raw score (13,14).

The hanging leg tuck (LTK) was started after a maximum four-minute rest from the previous event. Soldiers were allotted a total of two minutes to complete the event. A Soldier began by gripping the bar with and alternating grip, hands placed within six inches of each other. The body must hang fully extended. In the first phase, the Soldier pulled themselves up by flexing the knees, hips, waist and biceps until both knees touched both elbows. The second phase is controlled extension, returning to and resting in the starting point. Swinging to gain momentum was not allowed. Resting was only allowed while hanging. If a Soldier touched the ground to rest the event was terminated. The number of correctly performed leg tucks was recorded as the raw score. Safety was insured by maintaining a clean, dry bar, and monitoring Soldiers as they mounted and dismounted the bar (13,14).

The two mile run (2MR) was the final event and was started after a five minute maximum rest from the previous event. The run was conducted on a regulation sized rubberized track and timed by a pair of stop watches. Times were recorded to the nearest whole second (13,14).

Height was measured using a Height-Roller™ Wall Mounted Measuring Tape or similar approved tape measure. Soldiers stood with their heels against the wall without shoes, and their head level as determined by the square tape. Height was recorded to the nearest .5 inch (1.27 cm) on the DA Form 5500 in accordance to Army Regulation 600-9 (12).

Weight was measured using a calibrated Rubbermaid^®^ Digital Utility Scale, or similar with a 400 lbs. (181.4 kg) capacity measured in .5 lb. (0.22 kg) increments. Soldiers were weighed in standard PT uniform, without shoes. Weight was rounded down to the nearest full pound and recorded on the DA Form 5500 in accordance to Army regulation 600-9 (12). Any Soldier who exceeded the Army HT/WT standard was “taped” to determine the Soldier’s body composition.

A 0.25-inch-wide x 50-inch-long (6.35 mm x 127 cm) flexible measuring tape was to determine Army body composition. Neck circumference measurements were taken by placing the tape directly over the laryngeal prominence and circling the neck with the measuring tape parallel to the ground. Waist measurements were recorded using the same technique with the tape placed over the naval (Figure 2). Three measurements were taken at each location to mitigate error. The difference between the neck and waist measurements were recorded as the circumference value on the DA Form 5500 as directed by Army Regulation 600-9 (12). Because of the universal recognition, BMI was calculated and used for this study.

**Figure 2.**
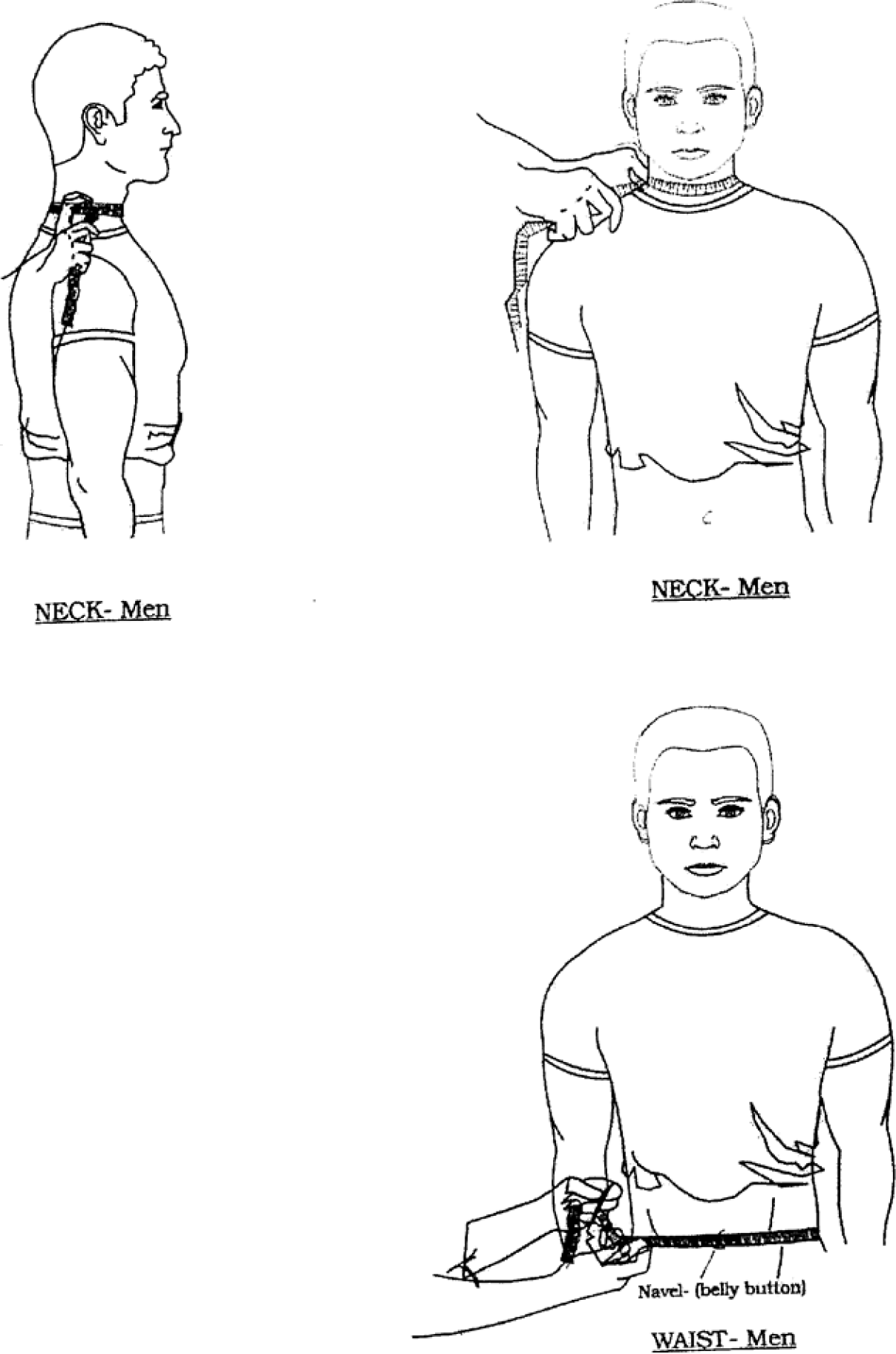
Male taping procedure. Diagram not to scale. (12) (Figure reprinted with permission of the U.S. Army).

### Statistical Analysis

The mean and standard deviation (SD) were calculated for the Soldier’s demographic information. The mean and SD were calculated for the ACFT event and total scores. Body mass index was calculated as body mass (kgs)/ (height (m)^2^). Pearson correlation coefficients (r) were calculated between the ACFT event scores, ACFT total, and BMI. Significance was considered as alpha≤0.05 when an |ES|≥0.40 was identified. An independent t-test was used to compare the ACFT scores between the Soldiers who passed the HT/WT standard and those how did not pass (α≤0.05). Analysis and data management were conducted with MS Excel 2013. The Excel spreadsheet was peer reviewed for exactitude as recommended by AlTarawneh and Thorne (1).

## RESULTS

Descriptive information of the participant Soldiers are presented in Table 1 (*n=*655). Table 2 provides the mean and standard deviation for the U.S. Army ACFT events and ACFT total score. Table 3 displays the correlations between the U.S. Army ACFT events, total, and Soldier BMI. Table 4 provides the Soldier’s height, weight, BMI and ACFT points for those who passed the HT/WT standard and those who did not pass. Likewise, Table 4 provides the results of a two-tailed t-test comparing BMI and ACFT scores between those who passed the HT/WT standard and those who did not pass.

**Table 1.** Descriptive Information

| Soldiers<br>Active Duty | Height (m) | Mass (kg) | BMI (kg/m<sup>2</sup>) |
| --- | --- | --- | --- |
| Males<br>(n=655) | 1.79±0.07 | 86.8±14.2 | 27.1±3.9 |

mean±standard deviation

**Table 2.** ACFT Test Scores

| MDL<br>(kgs) | SPT<br>(meters) | HRP<br>(reps) | SDC<br>(secs) | LTK<br>(reps) | 2MR<br>(secs) | ACFT<br>Total Score |
| --- | --- | --- | --- | --- | --- | --- |
| 92.2±31.8 | 9.5±2.2 | 24.6±13.1 | 119.8±21.7 | 6.2±5.4 | 1095.0±233.7 | 442.3±54.4 |

mean±standard deviation, MDL-3 rep max dead lift, SPT-standing power throw, HRP-hand release pushup, SDC-spring-drag-carry shuttle, LTK-hanging leg tuck, 2MR-two mile run, ACFT passing score $\geq 360$ points.

**Table 3.** ACFT Event Correlation Matrix and BMI

|  | MDL<br>(kgs) | SPT<br>(meters) | HRP<br>(reps) | SDC<br>(secs) | LTK<br>(reps) | 2MR<br>(secs) | ACFT<br>Total<br>Score |
| --- | --- | --- | --- | --- | --- | --- | --- |
| 3-RM DL | 1.00 |  |  |  |  |  |  |
| SPT | 0.45* | 1.00 |  |  |  |  |  |
| Push-up | 0.64* | 0.29 | 1.00 |  |  |  |  |
| SDC | -0.39 | -0.40* | -0.40* | 1.00 |  |  |  |
| Leg Tuck | 0.50* | 0.27 | 0.72* | -0.40* | 1.00 |  |  |
| Two Mile<br>Run | -0.11 | -0.02 | -0.22 | 0.32 | -0.31 | 1.00 |  |
| ACFT<br>Total<br>Score | 0.70* | 0.50* | 0.74* | -0.68* | 0.76* | -0.53* | 1.00 |
| BMI | 0.21 | 0.25 | -0.04 | 0.07 | -0.20 | 0.28 | -0.09 |

\*p-value<0.05 and |ES|≥0.40, MDL-3 rep max dead lift, SPT-standing power throw, HRP-hand release pushup, SDC-spring-drag-carry shuttle, LTK-hanging leg tuck, 2MR-two mile run.

**Table 4.**
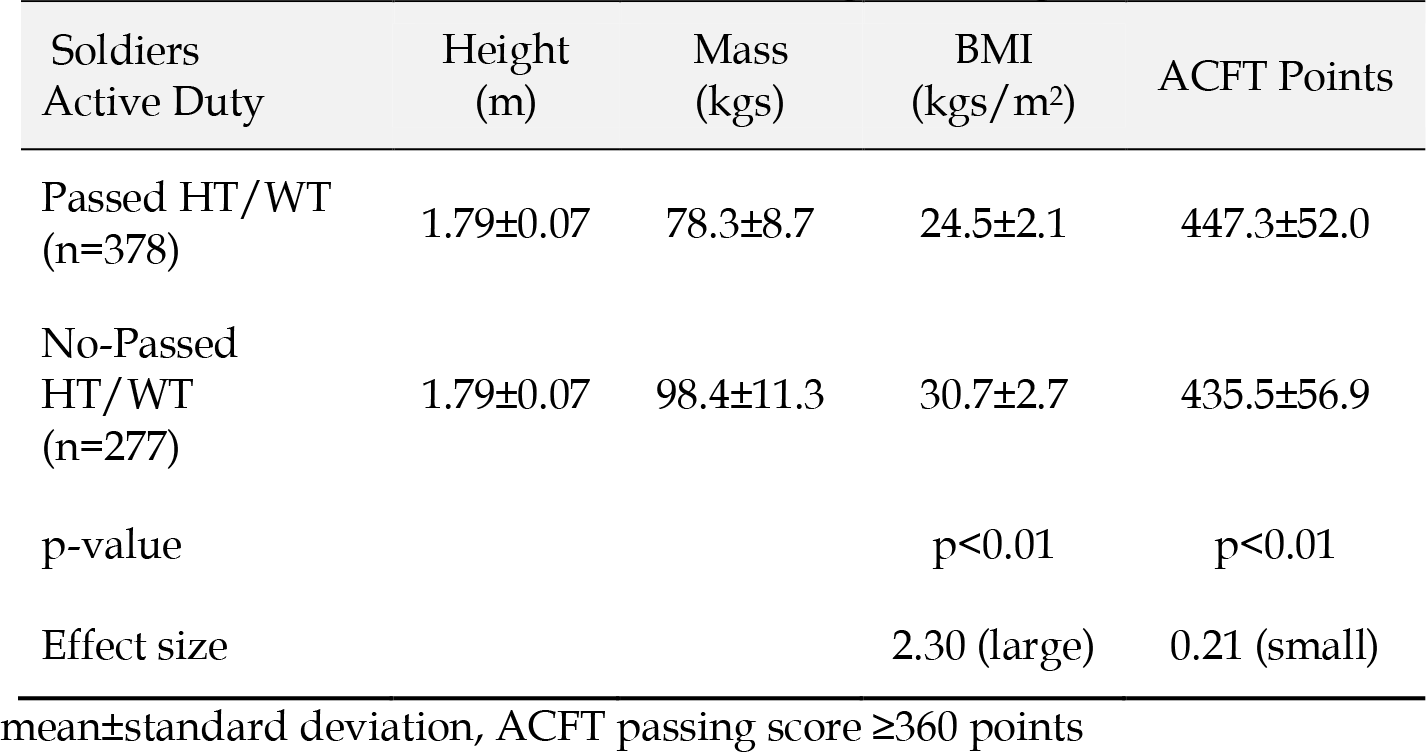
ACFT Scores Passed vs. No-Pass Height to Weight

| Soldiers<br>Active Duty | Height<br>(m) | Mass<br>(kgs) | BMI<br>(kgs/m<sup>2</sup>) | ACFT Points |
| --- | --- | --- | --- | --- |
| Passed HT/WT<br>(n=378) | 1.79±0.07 | 78.3±8.7 | 24.5±2.1 | 447.3±52.0 |
| No-Passed<br>HT/WT<br>(n=277) | 1.79±0.07 | 98.4±11.3 | 30.7±2.7 | 435.5±56.9 |
| p-value |  |  | p<0.01 | p<0.01 |
| Effect size |  |  | 2.30 (large) | 0.21 (small) |

mean±standard deviation, ACFT passing score ≥360 points

## DISCUSSION

The purpose of this study was to investigate a relationship between Soldier height, weight, body composition (BMI) and performance on the ACFT events and total score within the ranks of U.S. Army National Guard Soldiers between the dates of 06/30/2021 to 09/30/2021. Likewise, it was of interest to determine if the Soldiers who did not pass the HT/WT standard performed differently on the ACFT than the Soldiers who did pass the HT/WT standard. The results of the study indicated that correlations between the ACFT scores and BMI are weak. However, there was a significant difference in ACFT scores between those Soldiers who passed the HT/WT standard compared to those who did not, noting a small effect size to be discussed further below.

Each event of the ACFT has minimum and maximum scores as follows: MDL 63.6kg-154.5 kg (140-340lbs), SPT 4.5m-12.5m, HRP 10-60 repetitions, SDC 180 sec-93 sec (3:00-1:33 min), LTK 1-20 repetitions, and 2MR 1260 sec-810 sec (21:00-13:30 min). The minimum passing composite score is 360 points with a required minimum 60 points per event. There is a maximum of 100 points per event for 600 total. The results of the study determined the mean ACFT total score was 442.3±54.4 (Table 2) or 73.7% average per event, well above the 360 minimum passing score. It can be concluded that U.S. Army National Guard Soldiers in the 65^th^ Field Artillery Brigade are able to perform above the Army minimum standard. See Figure 3 in Appendix A for further ACFT scoring standards (14).

The results of this study demonstrated that of the 655 participants who completed all 6 events, only n=73 (11.2%) failed to meet the minimum standard for the ACFT by failing one individual event. The two-mile run accounted for n=69 of those who failed exceeding the 21:00 min minimum. The sprint-drag-carry event accounted for three Soldiers. Only one failed the hand release push-up event completing only six of the ten required push-ups.

The weak relationships between the ACFT scores and BMI suggests that body height and weight have little bearing on ACFT performance among these Soldiers. Interestingly, there was no difference in height of the Soldiers who passed or failed HT/WT standard. However, Soldiers who failed the HT/WT standard presented a mean increase in body mass of 20.1kg (44.3 lbs.). A supporting study of ROTC cadets performing the APFT (16) found no correlation between BMI and APFT scores. Likewise, a similar study of Army ROTC cadets determined no correlation between APFT scores and body fat percent measurements of any method (35).

The results of the study also demonstrated that 277 of the 655 (42.3%) Soldiers exceeded the Army Body Composition HT/WT standard. This standard is a scale that determines that for x height in inches a Soldier can weigh no more than y lbs. Soldiers who fail to meet the HT/WT standard are “taped” to determine the Soldier’s body composition (%BF). Of the 277 Soldiers who were “taped”, only 60 (9.1%) failed the %BF standard as assessed by the tape test or “busted tape”. This may be an indication that ≈200 Soldiers are being unnecessarily “taped”. Because age data was not gathered, we cannot conclude the specific age demographic that is most likely to exceed Army body composition standards.

The results also suggested that the ACFT scores were significantly lower for Soldiers who failed the height weight standard (n=277) compared to those who did pass (n=378). However, this results needs to be viewed with perspective. The average ACFT scores for those who passed the HT/WT standard and those who did not, both exceeded the ACFT passing score of 360. Further, the effect size difference between the ACFT scores was small (ES=0.21). Finally, lets assume the reliability coefficient for the ACFT was extremely high r=0.95. The smallest detectable difference would be SDD≈32 ACFT points (where: SDD=1.95*√2*SEM; SEM=SD√(1-0.95); SD=52.0). As such, the difference in mean AFCT scores between the Soldiers who failed the HT/WT standard compared to those who did pass is less than the SDD (i.e. within the range of error).

Regardless of their performance on the ACFT, Soldiers who fail Army body composition are barred from re-enlistment, promotion, attending military education and holding leadership positions. A common complaint among Army Soldiers is that the “tape test” is inaccurate and favors Soldiers who have a large neck and at least one study supports this argument (2). A Soldier with a large or even fat neck often has an advantage in relation to their waist circumference. For example: a 21-year-old male Soldier, 72 inches (182.9 cm) tall and weighing 250 lbs. (113.4kg) with a neck circumference of 20 inches (50.8 cm) and waist circumference of 42 inches (106.7kg) is determined to have 22% body fat and is in compliance with the Army standard. Calculated BMI for this Soldier is 33.9 and well in to the obese category. By contrast, a 21-year-old male Soldier, 72 inches (182.9 cm) tall and weighing 200 lbs. (90.7kg) with a neck circumference of 16 inches (40.6 cm) and waist circumference of 38.5 inches (97.8kg) is determined to have 23% body fat and exceeds the Army standard for his age. Calculated BMI for this Soldier is 27.1 and in the middle of the overweight category. Despite being smaller by all measures the latter fails the Army body composition standard.

The Army recently announced that a study would be conducted to determine a better way of measuring a Soldier’s body composition. The study will include: 3D full-body surface scan, DEXA scan, bioelectrical impedance, and the traditional tape test (19). The results of the study have the potential to more accurately assess a Soldier’s body composition. What is yet to be determined is if the Army will adjust the standard to reduce the number of Soldiers requiring body composition measurement particularly for those who successfully pass the ACFT.

The major assumption of this study is that Soldiers performed the ACFT with maximal effort. This early testing is considered “diagnostic”, recorded only as a practice test, and Soldiers potentially only preformed to the minimum required standard to pass the assessment. With that said, the average AFCT score of the Soldiers far exceeded the minimal AFCT passing score. A follow-on study using “For Record” data from the same population is merited as it may produce different results, presumably higher scoring then the current study. Administration of the ACFT is assumed to be conducted to the same standard across the Brigade which is comprised of 12 Company/Battery sized elements, in accordance with Army Regulation (12,13,14) and that all of the methods used in gathering data are reliable and sensitive. Specifically, that the same instrumentation and objectivity of grading is utilized with all non-timed events. Historically, the military is trained, funded and equipped to be standardized across all ranks and because of this, is able to conduct research with large samples in a controlled environment. For this study, the ACFT was conducted outdoors and weather could have impacted performance outcomes of the events. Testing was conducted from various locations on different days and times of the day. Temperature could have also affected the outcome of the Soldiers performance.

In summary, 88.9% (n=582) of the Field Artillery Soldiers tested met or exceed the minimum Army standard for combat fitness as assessed by the ACFT. Only 9.2% failed to meet the Army Body Composition standard by exceeding the authorized %BF as determined by the tape test. The study results indicated that little to no correlation existed between Soldier BMI and ACFT performance supporting the aforementioned findings (16, 35) and indicating that body composition (as assessed herein) is not a good predictor of ACFT performance. To attempt to answer our original questions; the results of this study suggest that the ACFT does not appear to favor larger or smaller Soldiers. The results also indicate that it may be time for the Army to reevaluate the current body composition standard as large Solder’s appear to be meeting the Army’s physical standard requirement. Future research should analyze the relationship of BMI and ACFT performance among female Soldiers to determine if similar results are found in the female population.

## ACKNOWLEDGEMENTS

The authors acknowledge the Army Human Research Protections Office (AHRPO) for approval to use of Soldier ACFT and HT/WT data. We also gratefully acknowledge the 65^th^ Field Artillery Brigade Command for providing Soldier data and supporting the study.

## Appendix A

**Figure 3.**
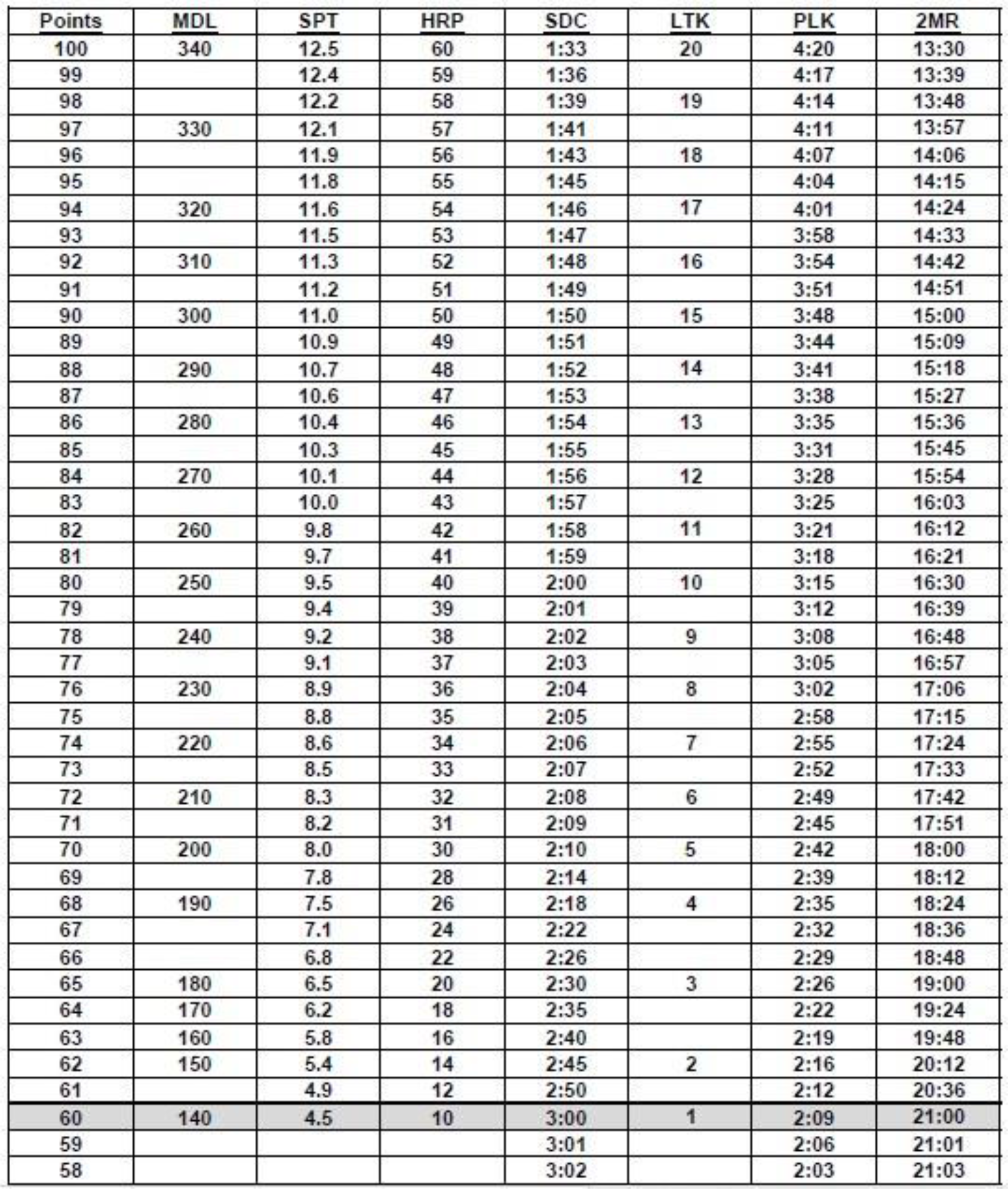
ACFT scoring standards. Minimum passing score of 60 points in each event. (Figure reprinted with permission of the U.S. Army). (14)

